# Use of a graph neural network to the weighted gene co-expression network analysis of Korean native cattle

**DOI:** 10.1101/2021.10.06.463300

**Authors:** Hyo-Jun Lee, Yoonji Chung, Ki Yong Chung, Young-Kuk Kim, Jun Heon Lee, Yeong Jun Koh, Seung Hwan Lee

## Abstract

In the general framework of the weighted gene co-expression network analysis (WGCNA), a hierarchical clustering algorithm is commonly used to module definition. However, hierarchical clustering depends strongly on the topological overlap measure. In other words, this algorithm may assign two genes with low topological overlap to different modules even though their expression patterns are similar. Here, a novel gene module clustering algorithm for WGCNA is proposed. We develop a gene module clustering network (gmcNet), which simultaneously addresses single-level expression and topological overlap measure. The proposed gmcNet includes a “co-expression pattern recognizer” (CEPR) and “module classifier”. The CEPR incorporates expression features of single genes into the topological features of co-expressed ones. Given this CEPR-embedded feature, the module classifier computes module assignment probabilities. We validated gmcNet performance using 4,976 genes from 20 native Korean cattle. We observed that the CEPR generates more robust features than single-level expression or topological overlap measure. Given the CEPR-embedded feature, gmcNet achieved the best performance in terms of modularity (0.261) and the differentially expressed signal (27.739) compared with other clustering methods tested. Furthermore, gmcNet detected some interesting biological functionalities for carcass weight, backfat thickness, intramuscular fat, and beef tenderness of Korean native cattle. Therefore, gmcNet is a useful framework for WGCNA module clustering.

**Author summary:** A graph neural network is a good alternative algorithm for WGCNA module clustering. Even though the graph-based learning methods have been widely applied in bioinformatics, most studies on WGCNA did not use graph neural network for module clustering. In addition, existing methods depend on topological overlap measure of gene pairs. This can degrade similarity of expression not only between modules, but also within module. On the other hand, the proposed gmcNet, which works similar to message-passing operation of graph neural network, simultaneously addresses single-level expression and topological overlap measure. We observed the higher performance of gmcNet comparing to existing methods for WGCNA module clustering. To adopt gmcNet as clustering algorithm of WGCNA, it remains future research issues to add noise filtering and optimal *k* search on gmcNet. This further research will extend our proposed method to be a useful module clustering algorithm in WGCNA. Furthermore, our findings will be of interest to computational biologists since the studies using graph neural networks to WGCNA are still rare.

## Introduction

Weighted gene co-expression network analysis (WGCNA) is often used to explore the system-level functionality of gene sets. WGCNA groups thousands of genes into a number of modules, simplifying biological interpretation. The general framework of a WGCNA [1] can be summarized as follows. First, the adjacencies of paired genes are calculated to define the gene co-expression network. The adjacencies are then incorporated into a topological overlap measure (TOM) to reveal gene-gene connections. Using the TOM, a clustering algorithm assigns intensively connected genes to the same modules. Finally, functional analyses are used to determine the biological meanings of the modules. This pipeline has been widely used in various fields. For example, recent biomedical studies used WGCNA to identify specific modules and hub genes related to human cancer [2] and arterial disease [3]. In animal and plant sciences, WGCNA has often been used to profile plant gene expression [4] and detect pathways responsible for complex animal traits [5, 6]. The module definitions greatly affect the interpretations of the results. WGCNA commonly uses a hierarchical clustering (HC) algorithm. This unsupervised clustering method places adjacent genes into the same modules based on pairwise TOM data. However, a concern has been raised that transformations of gene expressions into a TOM results in loss of raw-level expression features. HC-based module assignment depends strongly on the TOM. This can degrade similarity of expression not only between modules, but also within modules. In other words, HC may assign two genes with low topological overlap to different modules even though their expression patterns are similar. Furthermore, once a gene is added to a specific module, HC can never reverse the decision. This poses challenges when clustering complicated networks with many interconnected gene pairs. Thus, a new algorithm is needed to more accurately identify WGCNA gene modules. Langfelder et al. [7] developed a “dynamic tree cut” technique that clusters gene modules based on the shapes of dendrogram branches, but this still depends on TOM. Botía et al. [8] employed a derivative of K-means processing to refine gene modules generated by standard HC. However, this algorithm requires more than four steps beginning with module clustering, centroid computation, distance measurements, and gene relocation. This complex pipeline requires significant computational time and is thus unsuitable for very large networks.

A graph neural network (GNN) [9, 10] is a good alternative algorithm for module clustering. Given the recent successes of GNN, graph-based learning methods have been widely applied in bioinformatics. To predict drug-target interactions, recent studies employed various graphical convolutional networks [11, 12]. For single-cell RNA-seq analysis, a GNN was used to model cell-cell relationships [13] and impute gene expression levels within single cells [14]. Yang et al. [15] developed a GNN that extracted protein features from graphical information. However, most studies on WGCNA did not use GNN for module clustering.

In this paper, we introduce a GNN-based clustering algorithm for WGCNA: the gene module clustering network (gmcNet). Our method clusters genes based on their co-expression topologies (genes in the same module should be strongly connected) and single-level expression (genes in the same module should exhibit similar expression patterns). The main innovation of gmcNet is incorporating the expression feature of single gene with co-expression feature of their neighbor genes. gmcNet includes a “co-expression pattern recognizer” (CEPR) and a module classifier. The CEPR has a message-passing (MP) operation similar to that of GraphSAGE [16], except that the topological overlap matrix [1] is used as the input rather than the adjacency matrix. Using the former matrix, CEPR defines weighted relationships, consistent with the objective of WGCNA. The module classifier assigns genes to various modules using the CEPR-embedded features. We tested gmcNet using RNA-seq data for native Korean cattle, and compared the performance to that of other clustering algorithms. As GNNs are not widely used for WGCNA, our findings will be of interest to computational biologists.

## Results

### Model performance

To validate gmcNet performance, it was compared to four baseline clustering algorithms including HC, K-means clustering, K-medoids clustering, and random clustering (Fig 1). We measured performance in terms of clustering strength and functional enrichment. We used graph modularity [17] to measure the clustering strength, and the differentially expressed module (DEM) signals to assess functional enrichment.

**Fig 1.**
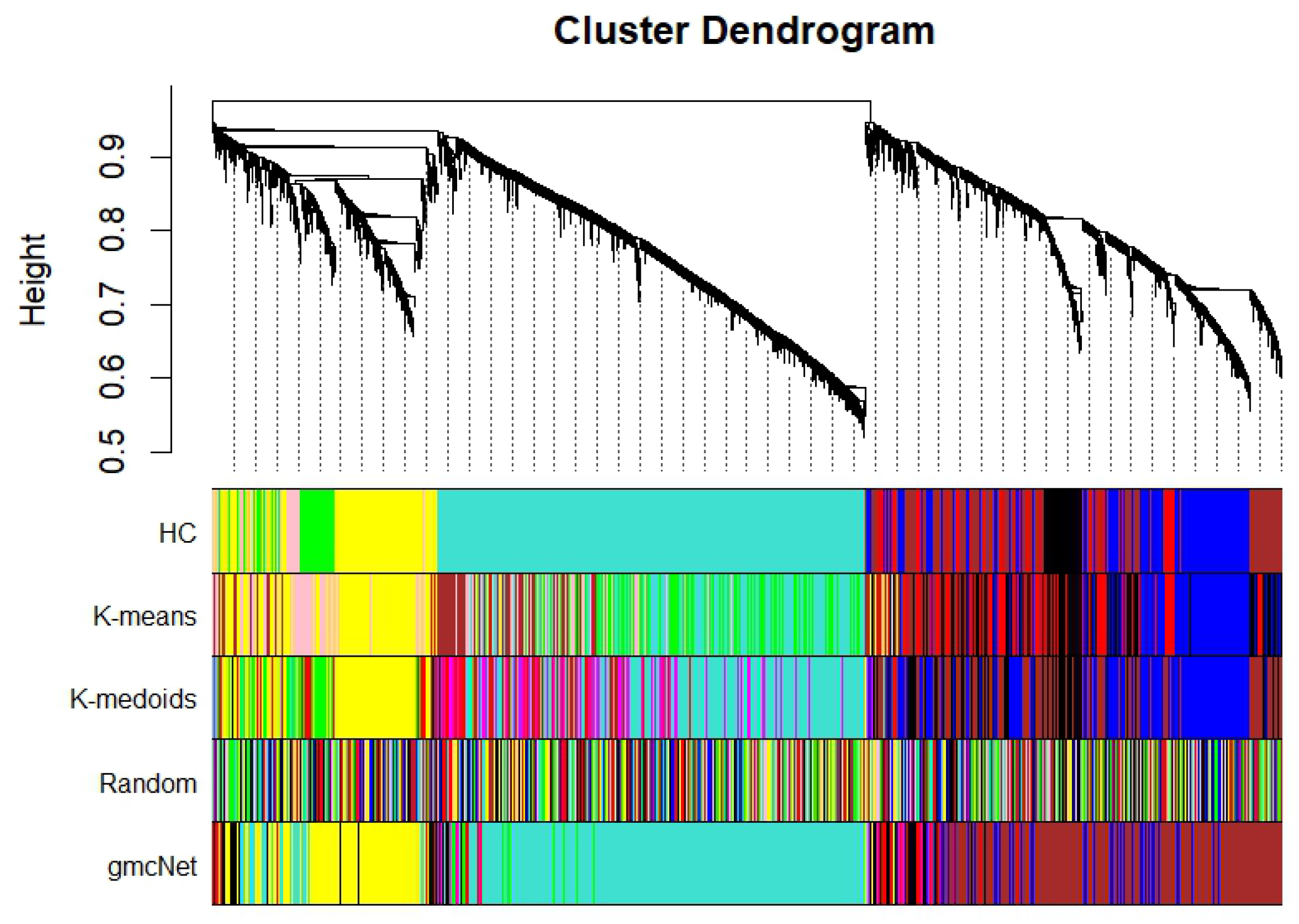
Module clustering results. The upper panel displays the hierarchical clustering dendrogram. In the lower panel, the colors show the module memberships determined by the methods on the left.

Table 1 presents the performances of the various methods. The single gene expression-based method (K-means) is robust to DEM signal capture, whereas the TOM-based methods (HC, K-medoids) provide higher modularity. On the other hand, gmcNet, which leverages both single gene expression and TOM, achieves the best DEM signal (27.739) and cluster modularity 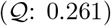. Comparison of gmcNet and HC revealed that gmcNet markedly increases modularity and the DEM signal by 0.042 and 9.121, respectively. Thus, gmcNet is more powerful than the other methods for revealing the apparent closeness of genes within the same module, and when making biological sense of the complex traits of native Korean cattle.

**Table 1.**
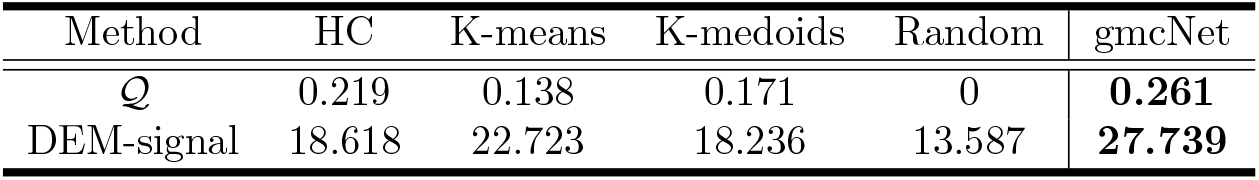
Model performance in terms of graph modularity 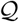 and DEM signaling.

| Method | HC | K-means | K-medoids | Random | gmcNet |
| --- | --- | --- | --- | --- | --- |
| $Q$ | 0.219 | 0.138 | 0.171 | 0 | **0.261** |
| DEM-signal | 18.618 | 22.723 | 18.236 | 13.587 | **27.739** |

### CEPR embedding

Fig 2 shows plots based on the first and second principal components of three feature types (single-level expression, TOM, and CEPR embedding). Single-level expression fails to distinguish modules with ambiguous boundaries. This may reflect the low modularity of K-means, which uses single-level expression for clustering. The TOM provides stronger connections between genes than single-level expression. However, it also decreases the distances between different modules and genes. As shown in Fig 2, K-medoids and HC, which use the TOM for clustering, do not clearly assign genes into different but closely related modules. Compared to the other types, CEPR embedding provides better separation, i.e. smaller distances between genes and larger ones between modules. With CEPR embedding, gmcNet defines gene modules more clearly and increases modularity.

**Fig 2.**
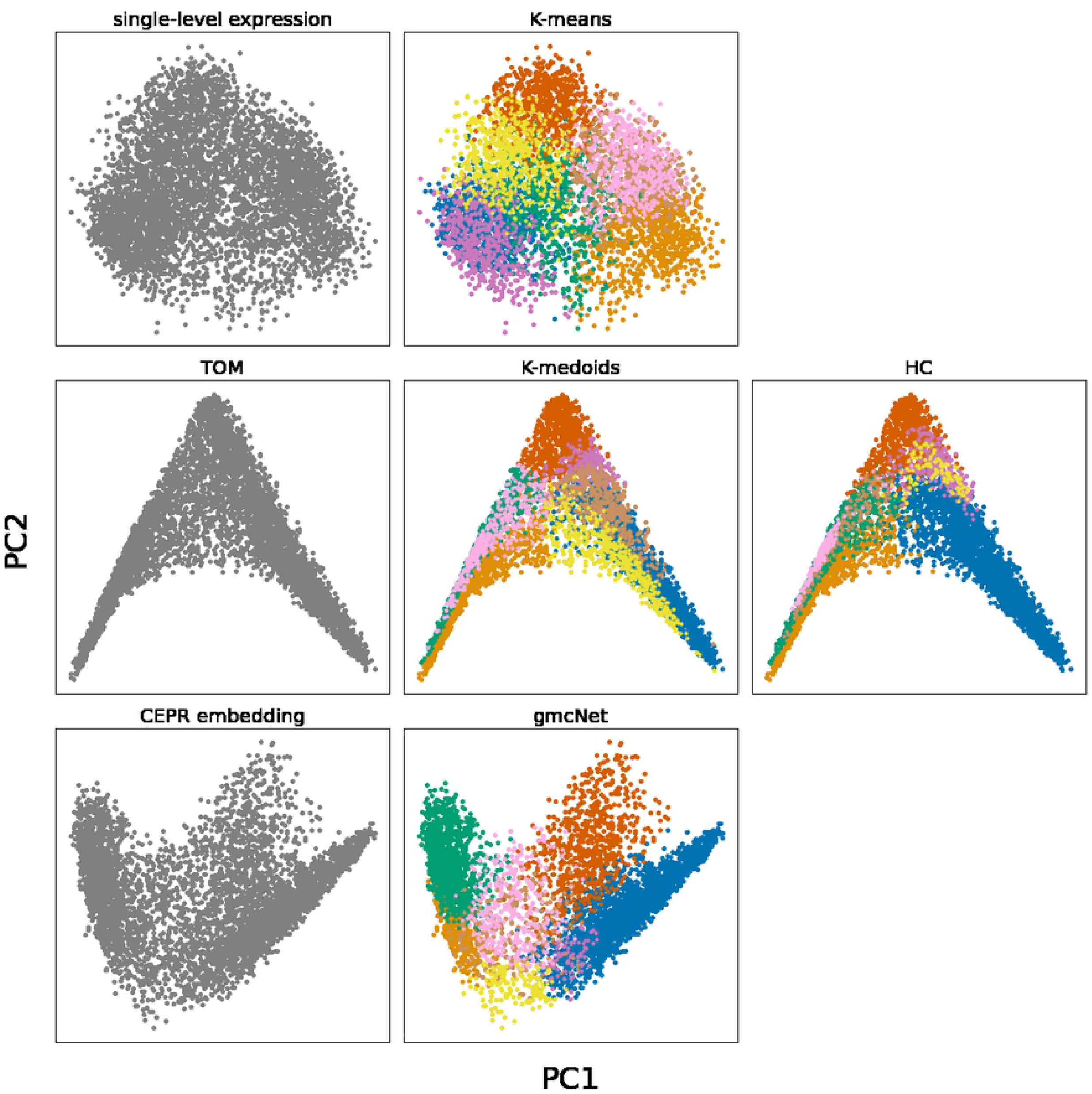
First and second principal components of three feature type and clustering results of each method. The x-axis and y-axis are first and second principal component. The colors show the module memberships determined by the methods on the top.

### Model performance at different *k* (number of clusters)

Our current implementation of gmcNet requires the setting of an optimal *k* (number of clusters). The effects of the *k*-value on modularity 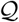 and the DEM signal are summarized in Fig 3. With an increasing *k*-value, the DEM signal increases while the 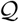 decreases. In contrast, gmcNet yields a larger DEM signal than HC even at smaller *k*-values (6 ≤ *k* < 8), and remains higher 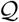 at larger *k*-values (*k* = 9). gmcNet outperforms K-means and K-medoids for all *k*-values. These results can demonstrate the superiority of gmcNet regardless of the *k*-value.

**Fig 3.**
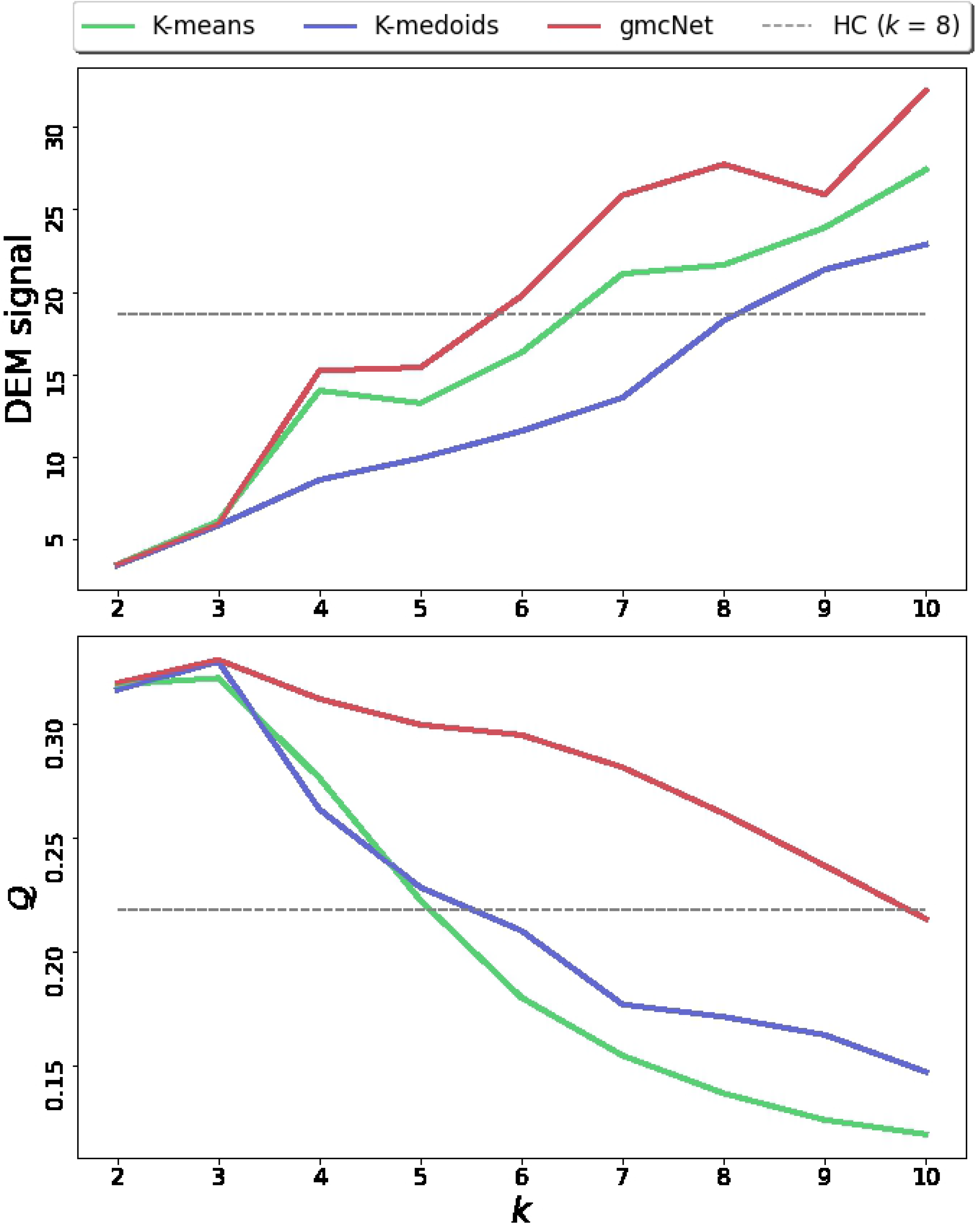
Optimal *k* searching considering DEM signaling and the modularity 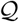.

### Functional enrichment analysis of native Korean cattle

To identify the DEMs, we performed linear regression analysis of the module eigengenes [1] for four complex traits, including carcass weight (CWT), backfat thickness (BF), intramuscular fat content (IMF), and the Warner-Bratzler shear force (WBSF). Fig 4 shows the results. In terms of the number of DEMs, IMF ranked first with four modules (K2, K3, K4, and K8) followed by BF (K2, K3, and K4), WBSF (K5 and K7), and CWT (K1 and K6). Interestingly, K5 and K7, which contain large numbers of genes, were significant to WBSF. This may reflect our mode of data collection; the RNA-seq data were from the *longissimus-dorsi* muscle and WBSF indicates the tenderness of beef muscle. Also, gmcNet detected 11 significant module-trait interactions. gmcNet found more DEMs than the other methods (HC: 9, K-means: 10, and K-medoids: 10) (S1 Fig).

**Fig 4.**
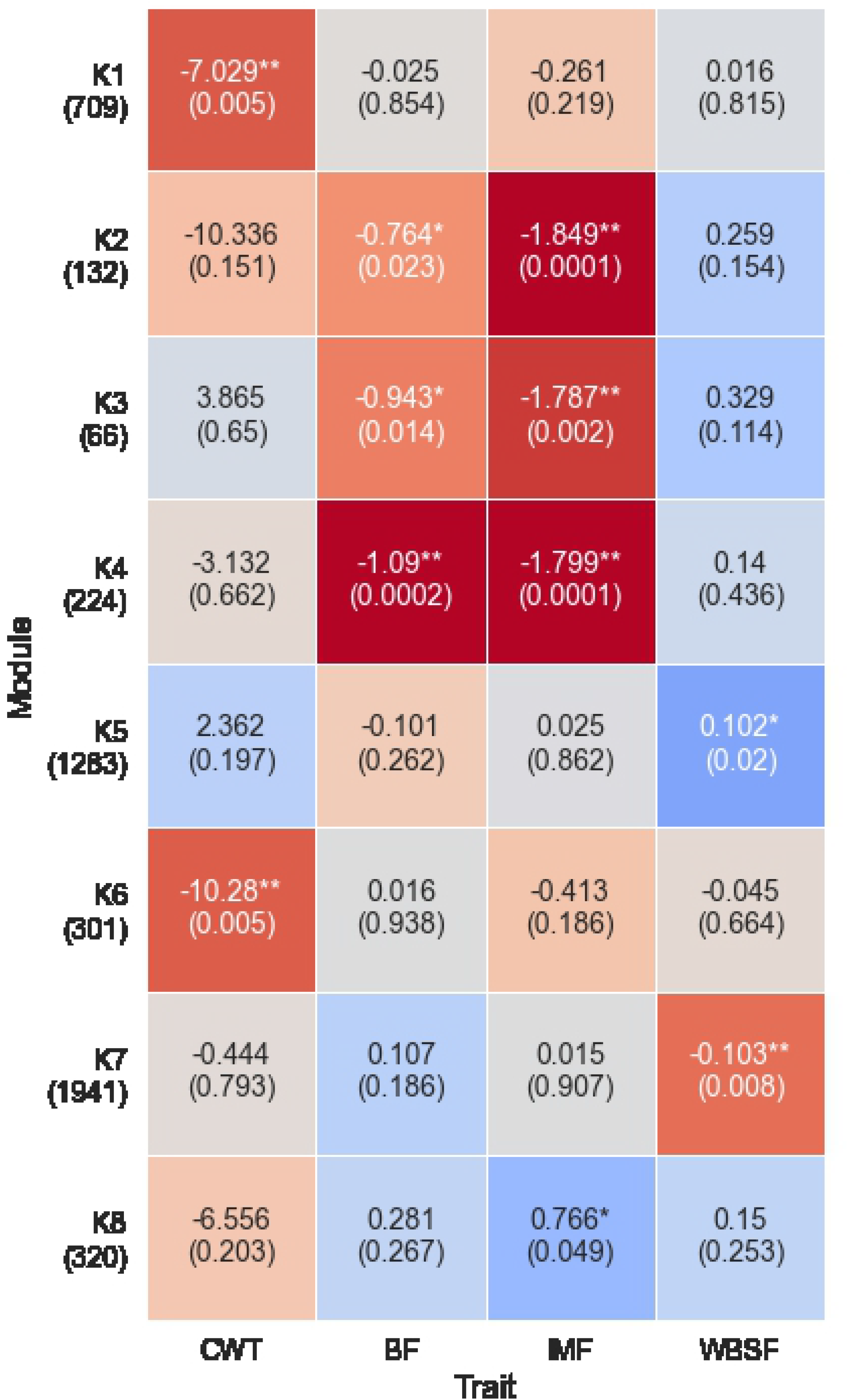
The DEM signals of modules defined by gmcNet. The y-axis shows the module names and numbers of genes within each module. The x-axis shows the complex traits. The numbers in each cell are regression coefficients (no parentheses) and the regression *p*-values (in parentheses). Red and blue indicate negative and positive coefficients, respectively. * *p* < 0.05, ** *p* < 0.01.

We used Gene Ontology (GO) enrichment analysis [18] to annotate the biological processes of the modules defined by gmcNet. Three modules (K1, K5, and K7) were linked to significant processes (Fig 5). K1, a CWT-related module, was enriched in “biosynthetic” and “metabolic” processes. Based on both the DEM analysis and the GO enrichment results, K1 seems to involve many genes associated with growth-related traits. Two WBSF-related modules (K5 and K7) were enriched in “immune system” and “protein catabolism”, respectively. Although several studies have suggested that the immune system plays a key role in cattle weight gain and feed efficiency [19, 20], the association between beef tenderness and immune pathways is a novel finding. Various studies have reported an association between “protein catabolic process” and beef tenderness [21–23]. Therefore, the results suggest that K7 is a key module of beef tenderness in native Korean cattle.

**Fig 5.**
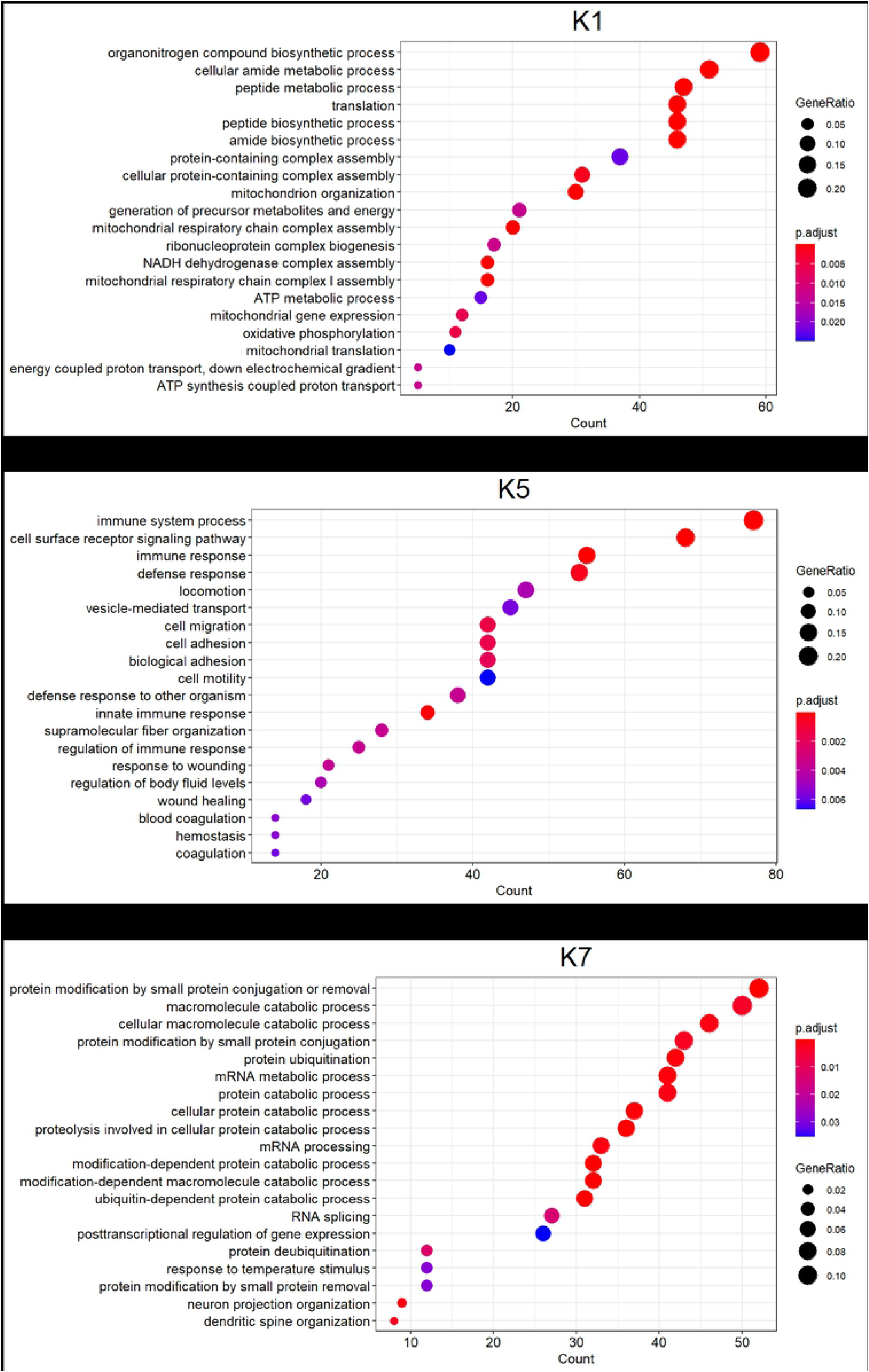
The biological processes of three significant modules: (a) K1, (b) K5, and (c) K7. p.adjust is a *p*-value adjusted by the Bonferroni method.

### Hub gene searches for modules of interest

Given the functional enrichment results, we selected the four modules, K1, K2, K4, and K7, as the principal modules of complex traits. Fig 6 shows the hub gene networks and Table 2 shows the related traits. The six hub genes of K1 are related to quantitative traits including growth (*LAMTOR5* [24] and *PAM16* [25]) and feed intake (*NDUFB1* [26], *NDUFB4* [27], *ATP5MF* [28], and *SEC61G* [29]). These findings support our suggestion that K1 is significant in terms of CWT. K2 and K4, associated with fat-related traits (BF and IMF) in DEM analysis, include eight (*ACSL3* [30], *NFKB1* [31], *CYP2R1* [32], *HSF2* [33], *TMEM135* [34], *PDCD4* [35], *HERPUD2* [36], and *NMRAL1* [37]) and seven (*SPNS1* [31], *MYOD1* [38], *PDXK* [39], *TMUB1* [34], *ARHGAP26* [40], *RAB15* [31], and *TP73* [41]) fat-related hub genes, respectively. Thus, future research should identify the relationships between fat metabolism and modules K2 and K4. Although K7 was associated with WBSF in DEM analysis, only four hub genes (*PARD3* [42], *EIF4G3* [43], *PAFAH1B1* [44], and *CAMTA2* [45]) were associated with growth-related traits; the other hub genes were all novel.

**Fig 6.**
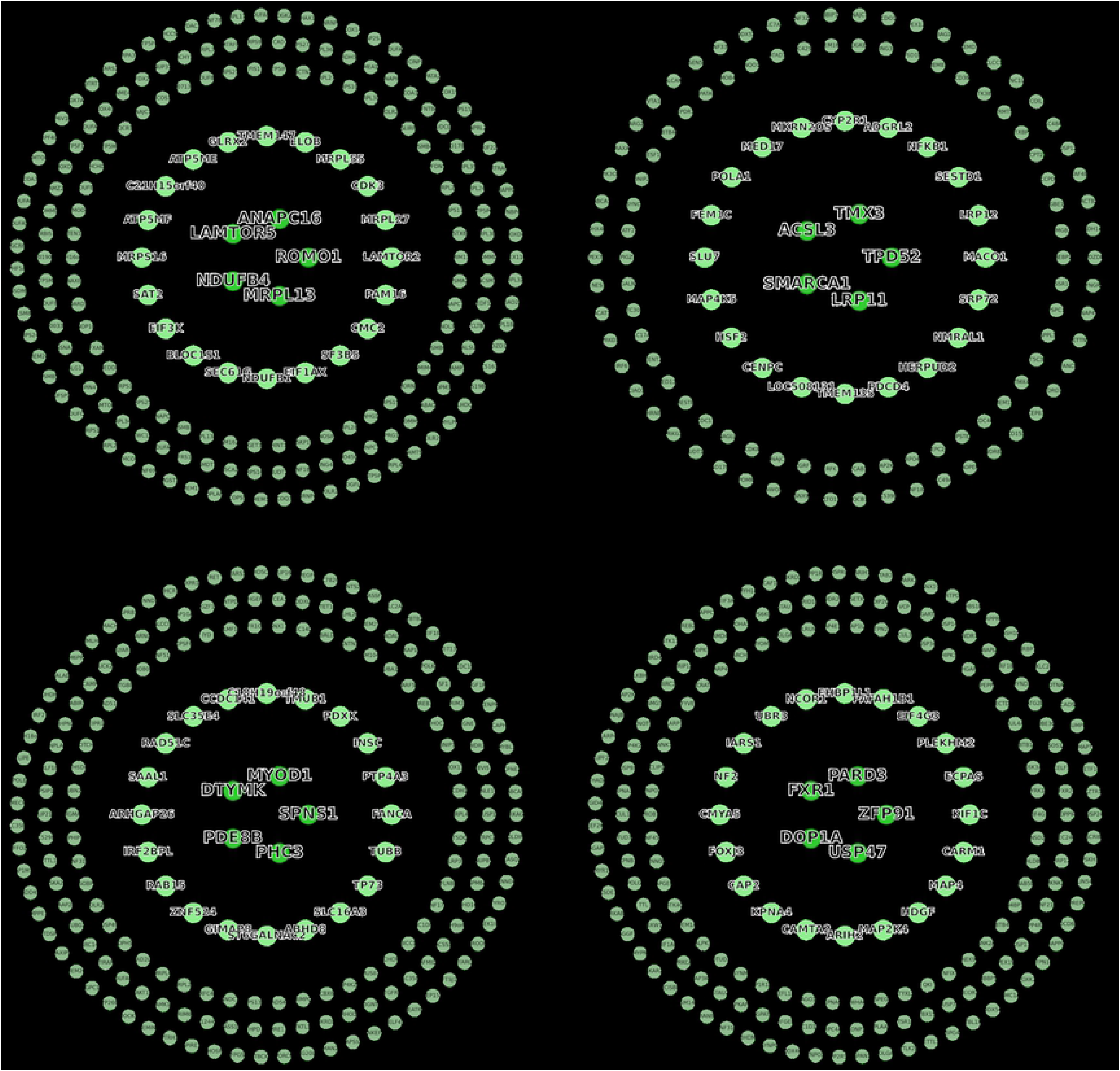
Hub gene networks of the four principal modules of native Korean cattle: (a) K1, (b) K2, (c) K4, (d) K7. From the outside in, the top 200, top 25, and top 5 hub genes are shown. The linkages of the top 5 hub genes are shown as the edges of the networks.

**Table 2.** Hub genes and associated traits of the main modules.

| Module | Hub gene<sup>1</sup> | Significant trait<sup>2</sup> | Reported cattle traits affected |
| --- | --- | --- | --- |
| K1 | *ROMO1, ANAPC16, LAMTOR5, NDUFB4, MRPL13, LAMTOR2, MRPL27, CDK3, MRPL55, ELOB, TMEM147, GLRX2, ATP5ME, C21H15orf40, ATP5MF, MRPS16, SAT2, EIF3K, BLOC1S1, SEC61G, NDUFB1, EIF1AX, SF3B5, CMC2, PAM16* | CWT** (0.005) | Growth [24, 25]; Tenderness [7, 46, 47, 47]; Feed intake [26, 28, 29]; Fat [31, 48, 49] |
| K2 | *TPD52, TMX3, ACSL3, SMARCA1, LRP11, MACO1, LRP12, SESTD1, NFKB1, ADGRL2, CYP2R1, MKRN2OS, MED17, POLA1, FEM1C, SLU7, MAP4K5, HSF2, CENPC, LOC508131, TMEM135, PDCD4, HERPUD2, NMRAL1, SRP72* | BF* (0.023); IMF** (0.0001) | Fat [30, 31, 50]; Growth [51–53] |
| K4 | *SPNS1, MYOD1, DTYMK, PDE8B, PHC3, FANCA, PTP4A3, INSC, PDXK, TMUB1, C18H19orf48, CCDC141, SLC35E4, RAD51C, SAAL1, ARHGAP26, IRF2BPL, RAB15, ZNF524, GIMAP8, ST6GALNAC2, ABHD8, SLC16A3, TP73, TUBB* | BF** (0.0002); IMF** (0.0001) | Fat [31, 38, 39]; Growth [48, 54, 55]; Feed intake [5, 56, 57]; Tenderness [58, 59] |
| K7 | *ZFP91, PARD3, FXR1, DOP1A, USP47, KIF1C, ECPAS, PLEKHM2, EIF4G3, PAFAH1B1, EHBP1L1, NCOR1, UBR3, IARS1, NF2, CMYA5, FOXJ3, CAP2, KPNA4, CAMTA2, ARIH2, MAP2K4, HDGF, MAP4, CARM1* | WBSF** (0.008) | Fat [60]; Growth [42–44] |

Hub gene<sup>1</sup>: Top 25 hub genes; Significant trait<sup>2</sup>: Significant traits revealed by DEM analysis (*p*-values). The reported traits affected by each hub gene are listed in S3 Table.

## Discussion

Single-level expression is generally appropriate to identify trait-specific marker genes that are differentially expressed depending on the biological phenotype [61]. Here, we found that single-level expression also revealed trait-specific modules with strong DEM signals. However, most existing WGCNA methods address only the co-expression topology (including TOM); the DEM signals are weak. On the other hand, our gmcNet simultaneously addresses single-level expression and TOM. gmcNet thus yielded larger DEM signals than other clustering methods. Furthermore, gmcNet produced some novel and interesting results. Threfore, gmcNet can detect module functionality and improves our understanding of WGCNA system-level biology. Also, gmcNet yields strong adjacencies between genes in the same module. gmcNet exploits the learnable properties of CEPR, which aggregates single-gene expressions with the co-expression features of its first neighbors, embedding these features to reduced dimensions. As noted in the Results section, CEPR generates more robust features than single-gene expression data or TOM. Given the CEPR-embedded feature, gmcNet achieved the best WGCNA modularity of all clustering methods tested.

Many genes are uniformly expressed in all individuals. Such genes (“noise”) are intimately connected with nested modules and exhibit no differential expression in complex trait analysis. Any attempt to cluster them disrupts module identification and obscures the biological implications. HC uses a dendrogram cut-off to exclude noisy genes. On the other hand, gmcNet assigns every gene to the most probable module. This may yield some meaningless assignments, because uniform expression may render the assignments to nested modules similar. Therefore, in future, it will be important to eliminate noise. We are exploring probability thresholding to this end. Specifically, genes with maximum probabilities lower than a given threshold will be excluded from module assignment. We will also add the optimal *k* search method to gmcNet; *k*-values can greatly increase model performance and may be modified depending on the characteristics of a dataset. Here, gmcNet used the optimal *k* of HC and performed better than other methods. In addition, gmcNet outperformed K-means and K-medoids at all *k*-values tested (2-10). Thus, the addition of an optimal *k* search would improve gmcNet performance in the context of WGCNA.

## Conclusion

We derived a gene module clustering network, gmcNet, which simultaneously addresses single-level expression and TOM. We validated gmcNet performance using 4,976 genes from 20 native Korean cattle. gmcNet reliably assigned genes to modules exhibiting high modularity and DEM signals. gmcNet also detected some interesting biological functionalities. Therefore, gmcNet is a useful framework for WGCNA module clustering.

## Materials and methods

### Data collection

A total of 20 native Korean steers were used; all were humanely slaughtered at 30 months of age. The CWT (kg), and BF (mm) were measured after chilling for 24 hours. BF was measured at the junction of the 12th and 13th ribs. The WBSF and IMF were measured at the *longissimus-dorsi* muscle according to [62] and [63], respectively. RNA from the *longissimus-dorsi* muscle was extracted using TRIzol reagent (Invitrogen, Carlsbad, CA, USA). RNA quality and quantity were assessed by automated capillary gel electrophoresis performed using a Bioanalyzer 2100 running the RNA 6000 Nano LabChip (Agilent Technologies Ireland, Dublin, Ireland). Only RNA samples with RNA integrity ≥ 7 were retained. Reads per kilobase per million (RPKM) were computed for each gene. After deriving correlation coefficients, we excluded 7,555 genes that exerted no effect (*p*-value > 0.1) on any of the four traits. Finally, 4,976 genes in 20 samples were subjected to this study. The National Institute of Animal Science (NIAS) of the Rural Development Administration (RDA) of South Korea approved the experimental procedures.

### Co-expression network construction

To represent the co-expression network in matrix form, we used the topological overlap matrix of [1]. Briefly, the adjacency of each pair of genes *i* and *j* is given by *a_ij_* = |*cor_ij_*|*^β^* where *β* is a smoothing parameter and *cor_ij_* is the correlation coefficient between the single-level expressions of the two genes. Given the adjacency values *a_ij_*, the topological overlap matrix 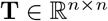 was created using a TOM [64], where *n* is the number of genes. Also, we constructed two additional topological overlap matrices to train gmcNet (Fig 7). 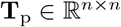, representing the positive network, was created leaving only positive correlation coefficients, whereas 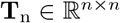, representing the negative network, was created leaving only negative correlation coefficients. After scale-free model fitting [1], we chose *β* = 6, *β* = 9, and *β* = 10 as the smoothing parameters for **T**, **T**_p_, and **T**_n_, respectively.

**Fig 7.**
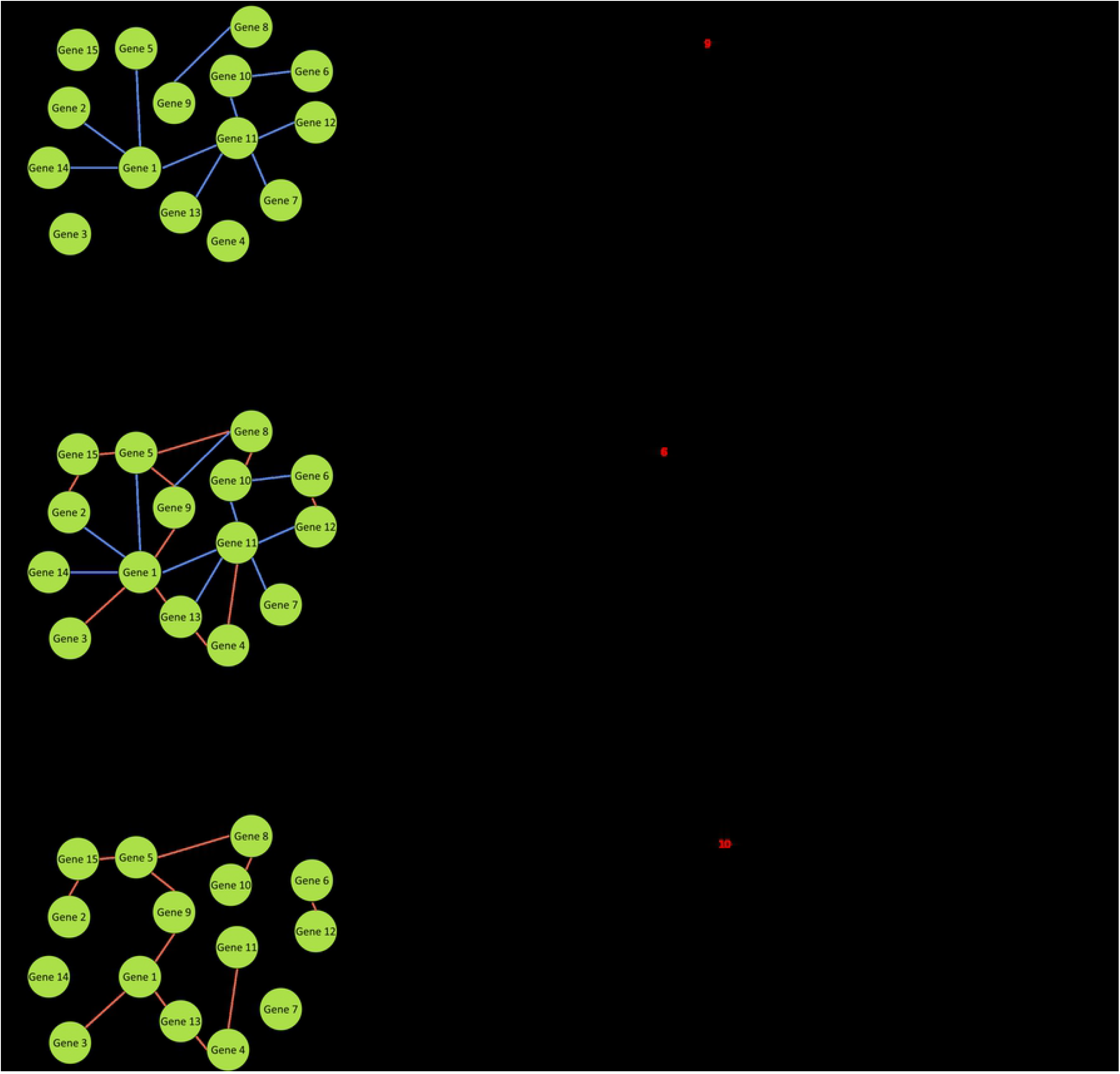
Construction of three topological overlap matrices. **T** is the topological overlap matrix of all relationships. **T**_p_ and **T**_n_ are the topological overlap matrices of positive and negative relationships respectively.

### Gene module clustering network

We developed a gene module clustering network (gmcNet) that clusters genes according to their co-expression topologies (genes in the same module should be strongly connected) and their single-level expression (genes in the same module should exhibit similar expression patterns). Fig 8 shows an overview of gmcNet, which features a co-expression pattern recognizer (CEPR) and module classifier. The CEPR incorporates the expression features of single genes into the topological features of co-expressed ones. Given this CEPR-embedded feature, the module classifier computes module assignment probabilities.

**Fig 8.**
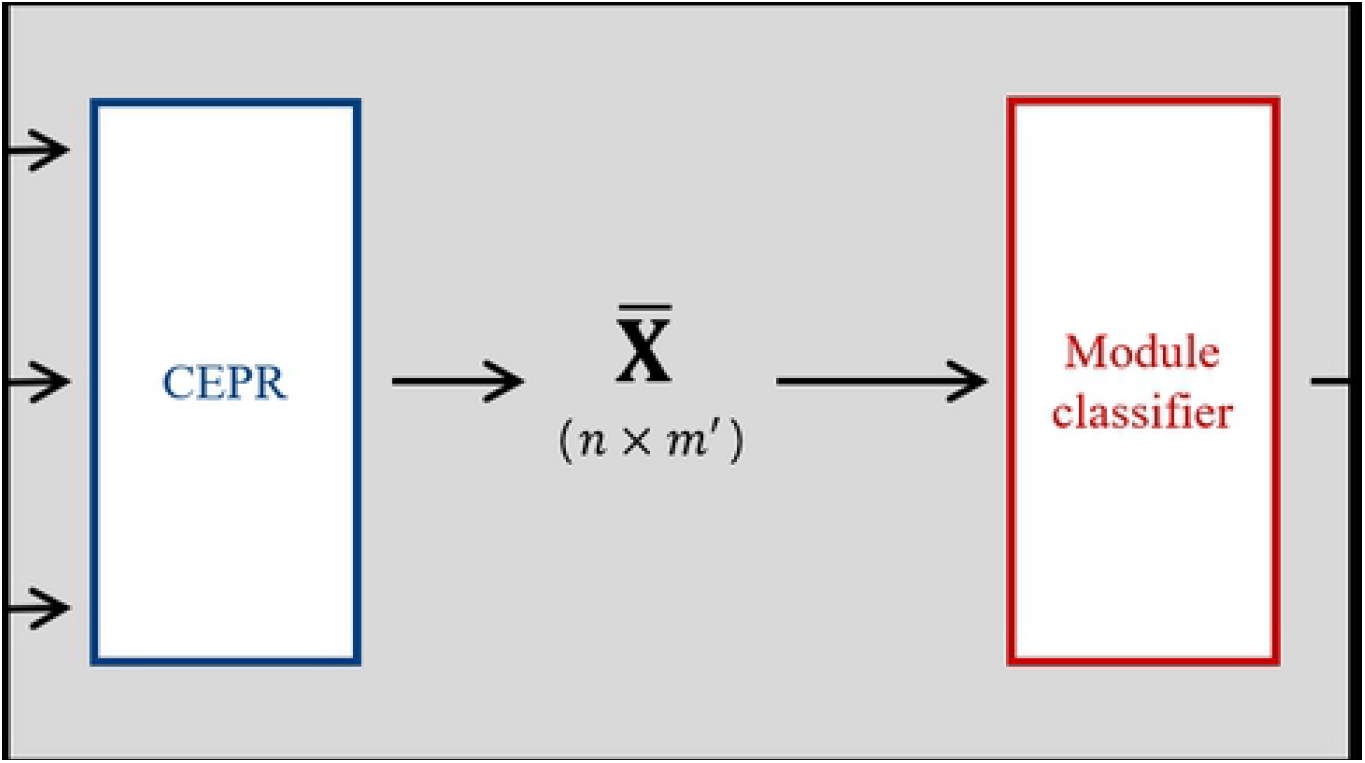
The architecture of gmcNet. 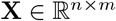 is the single-level expression of *n* genes in *m* samples. 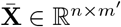 is CEPR-embedded feature with *m′* dimension. 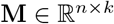 is assignment probability matrix of *n* genes to *k* modules. 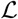 is loss function.

### Network structure

#### CEPR

The goal of CEPR is to integrate single-expression features with co-expression features. To achieve this, we used the MP operation of GraphSAGE [16], but employed the topological overlap matrix rather than the adjacency matrix. We computed a new topological overlap matrix 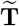 by zeroing the diagonal of **T** and applying degree normalization:

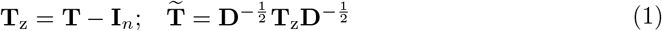

where **D** = diag(**T**_z_1*_n_*) is a degree matrix. Let 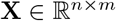 be the single-level expression of *n* genes in *m* samples. Then, single and co-expression can be simply combined via an MP operation:

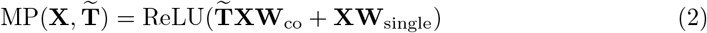

where **W**_co_ and **W**_single_ are the trainable parameters of the co- and single-expression features. As 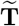 includes the topological adjacencies between gene pairs, it is easy to see that 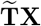 can be interpreted as a co-expression feature.

A simple MP operation cannot separate positive and negative co-expressions, even when they differ in different biological pathways. Therefore, we refined a simple MP to become a CEPR, as follows:

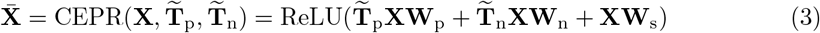

where 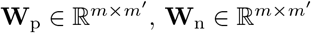, and 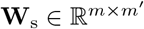 are the trainable weights of the positive co-expression, negative co-expressions, and single-expression, respectively. *m′* is an embedding dimension (set to 8). As 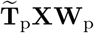 and 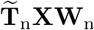 are identical in terms of dimensionality, CEPR learns various co-expressions by simply adding them. By skip connections of single-expression **XW**_s_, CEPR generates the embedding feature 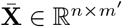, which deals with single-expression and two different co-expressions in the *m′* dimension.

#### Module classifier

Given the CEPR-embedded feature 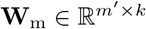, the module classifier computes a module assignment probability using a multi-layer perceptron (MLP):

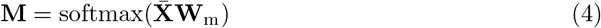

where 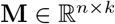 are the trainable weights for clustering of *k* modules. As softmax activation guarantees that *m_ij_* ∈ [0, 1], the *i*th-row of 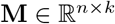 corresponds to the module-assignment probability of gene *i*. In other words, gene *i* belongs to module *c* if *m_ic_* is the maximum value of the *i*th-row of **M**.

#### Loss function

For unsupervised clustering, we employed the cut and orthogonality loss terms of MinCutPool [65]. The loss function when training gmcNet was defined as:

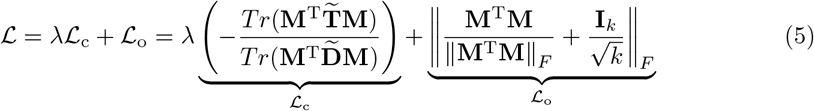

where ║·║*_F_* indicates the Frobenius norm and *T_r_* is the trace; λ is a balancing hyper-parameter, which is set to 2.6. The cut loss term, 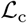, encourages clustering of strongly connected genes within the same module, and the orthogonality loss term, 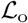, penalizes assignment to similarly sized modules.

#### Implementation Details

The model was iterated for 5,000 epochs using a GeForce RTX 2080ti. For the first 100 epochs, the balancing hyperparameter λ was set to 0 and the learning rate to 0.01. This prevented the creation of empty modules. After epoch 100, we set λ to 2.6 and the learning rate to 0.001. Model training was early stopped at 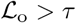, where *τ* is the orthogonal threshold, which was set to 0.8. The Adam optimizer [66] was used to minimize the loss function. Finally, **M** at the end of training was used for module assignment.

### Model performance

To validate gmcNet performance, HC [7], K-means [67] and K-medoids [68] were also used for module clustering and the results were compared to those of gmcNet. K-means uses single-expression feature **X** as input data; the HC and K-medoids use the topological distances 1 – **T** as inputs. The optimal *k* for K-means, K-medoids, and gmcNet was set to 8, as suggested by application of the dynamic tree cut technique [7] to HC.

#### Metrics

We measured the model performance in terms of modularity and DEM signaling. Module modularity is a commonly used metric in graph clustering. In a fully random graph, gene *i* and *j* of degrees *c_i_* = ∑*_u_ t_iu_* and *c_j_* = ∑*_u_ t_ju_* are connected with a probability *c_i_c_j_/s*, where *s* is the total topological overlap *s* = ∑*_ij_ t_ij_*. Modularity measures the divergence between intra-module connections as:

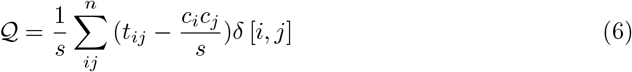

where *δ* [*i, j*] = 1 if *i* and *j* belong to the same module; otherwise, *δ* [*i, j*] = 0.

To measure DEM signals, we employed linear regression analysis to the module eigengenes, *i.e*. the first principal components of the modules, for four complex traits: CWT, BF, IMF and WBSF. Let *ρ*[*l, t*] = 1 if module *l* is significant (≤ 0.05) for trait t; otherwise, *ρ* [*l, t*] = 0. The final DEM signal was defined as:

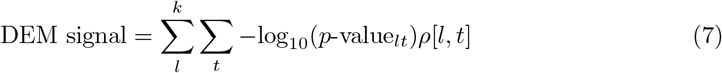

where *t* ∈ {CWT, BF, IMF, WBSF} and *p*-value*_lt_* indicates the significance value of module *l* in terms of trait *t*.

#### Functional enrichment analysis

The Bioconductor R package “clusterProfiler” [69] was used for GO analysis. The adjusted *p*-value (obtained using the Bonferroni method) was employed to examine the significance (p.adjust< 0.05) of all GO terms. The top 20 biological processes were extracted if there were more than 20 significant results. To identify hub genes, we calculated the correlation coefficients between single-level expression of each gene and the ME of the module it belong to. The top 25 genes (in terms of correlation coefficients) were defined as hub genes.

## Acknowledgments

This work was supported by Institute of Information & communications Technology Planning & Evaluation(IITP) grant funded by the Korea government(MSIT)(No.2020-0-01441, Artificial Intelligence Convergence Research Center(Chungnam National University)).

## Data availability

The gmcNet code and example data is available on GitHub at https://github.com/gywns6287/gmcNet. Request for full gene expression data of Korean native cattle can be made to Korea National Institute of Animal Science, Animal Genome & Bioinformatics Division (http://www.nias.go.kr/english/sub/boardHtml.do?boardId=depintro).

## Supporting information

**S1 Fig. The DEM signals of modules defined by baseline methods.**

**S2 Fig. Hub gene networks of the minor modules.**

**S3 Table. The reported traits affected by each hub gene.**

## Author Contributions

**Conceptualization:** Seung Hwan Lee, Yeong Jun Koh.

**Data Curation:** Ki Yong Chung.

**Formal Analysis:** Hyo-Jun Lee, Jun Heon Lee.

**Funding Acquisition:** Jun Heon Lee, Young-Kuk Kim.

**Methodology:** Hyo-Jun Lee, Young-Kuk Kim.

**Software:** Hyo-Jun Lee, Yeong Jun Koh.

**Visualization:** Yoonji Chung.

**Writing – Original Draft Preparation:** Hyo-Jun Lee, Yoonji Chung.

**Writing – Review & Editing:** Seung Hwan Lee, Yeong Jun Koh.

## Notes

### Competing Interest Statement

The authors have declared no competing interest.

## References

1. Zhang B, Horvath S. A general framework for weighted gene co-expression network analysis. Statistical applications in genetics and molecular biology. 2005;4(1).

2. Li J, Zhou D, Qiu W, Shi Y, Yang JJ, Chen S, et al. Application of weighted gene co-expression network analysis for data from paired design. Scientific reports. 2018;8(1):1–8.

3. Zheng PF, Chen LZ, Guan YZ, Liu P. Weighted gene co-expression network analysis identifies specific modules and hub genes related to coronary artery disease. Scientific Reports. 2021;11(1):1–13.

4. Rao X, Dixon RA. Co-expression networks for plant biology: why and how. Acta biochimica et biophysica Sinica. 2019;51(10):981–988.

5. Salleh M, Mazzoni G, Höglund J, Olijhoek D, Lund P, Løvendahl P, et al. RNA-Seq transcriptomics and pathway analyses reveal potential regulatory genes and molecular mechanisms in high-and low-residual feed intake in Nordic dairy cattle. BMC genomics. 2017;18(1):1–17.

6. Silva-Vignato B, Coutinho LL, Poleti MD, Cesar AS, Moncau CT, Regitano LC, et al. Gene co-expression networks associated with carcass traits reveal new pathways for muscle and fat deposition in Nelore cattle. BMC genomics. 2019;20(1):1–13.

7. Langfelder P, Zhang B, Horvath S. Defining clusters from a hierarchical cluster tree: the Dynamic Tree Cut package for R. Bioinformatics. 2008;24(5):719–720.

8. Botía JA, Vandrovcova J, Forabosco P, Guelfi S, D’Sa K, Hardy J, et al. An additional k-means clustering step improves the biological features of WGCNA gene co-expression networks. BMC systems biology. 2017;11(1):1–16.

9. Kipf TN, Welling M. Semi-supervised classification with graph convolutional networks. ICLR-17. 2017;.

10. Xu D, Zhu Y, Choy CB, Fei-Fei L. Scene graph generation by iterative message passing. In: Proceedings of the IEEE conference on computer vision and pattern recognition; 2017. p. 5410–5419.

11. Peng J, Wang Y, Guan J, Li J, Han R, Hao J, et al. An end-to-end heterogeneous graph representation learning-based framework for drug–target interaction prediction. Briefings in Bioinformatics. 2021;.

12. Zhao T, Hu Y, Valsdottir LR, Zang T, Peng J. Identifying drug–target interactions based on graph convolutional network and deep neural network. Briefings in bioinformatics. 2021;22(2):2141–2150.

13. Wang J, Ma A, Chang Y, Gong J, Jiang Y, Qi R, et al. scGNN is a novel graph neural network framework for single-cell RNA-Seq analyses. Nature communications. 2021;12(1):1–11.

14. Rao J, Zhou X, Lu Y, Zhao H, Yang Y. Imputing single-cell RNA-seq data by combining graph convolution and autoencoder neural networks. Iscience. 2021;24(5):102393.

15. Yang F, Fan K, Song D, Lin H. Graph-based prediction of Protein-protein interactions with attributed signed graph embedding. BMC bioinformatics. 2020;21(1):1–16.

16. Hamilton WL, Ying R, Leskovec J. Inductive representation learning on large graphs. In: Proceedings of the 31st International Conference on Neural Information Processing Systems; 2017. p. 1025–1035.

17. Newman ME. Modularity and community structure in networks. Proceedings of the national academy of sciences. 2006;103(23):8577–8582.

18. Wu T, Hu E, Xu S, Chen M, Guo P, Dai Z, et al. clusterProfiler 4.0: A universal enrichment tool for interpreting omics data. The Innovation. 2021; p. 100141.

19. Reynolds J, Foote A, Freetly H, Oliver W, Lindholm-Perry A. Relationships between inflammation-and immunity-related transcript abundance in the rumen and jejunum of beef steers with divergent average daily gain. Animal genetics. 2017;48(4):447–449.

20. Alexandre PA, Kogelman LJ, Santana MH, Passarelli D, Pulz LH, Fantinato-Neto P, et al. Liver transcriptomic networks reveal main biological processes associated with feed efficiency in beef cattle. BMC genomics. 2015;16(1):1–13.

21. Zhao C, Zan L, Wang Y, Updike MS, Liu G, Bequette BJ, et al. Functional proteomic and interactome analysis of proteins associated with beef tenderness in Angus cattle. Livestock Science. 2014;161:201–209.

22. Tian X, Wu W, Yu Q, Hou M, Jia F, Li X, et al. Quality and proteome changes of beef M. longissimus dorsi cooked using a water bath and ohmic heating process. Innovative Food Science & Emerging Technologies. 2016;34:259–266.

23. Li Y, Jin H, Yan C, Seo K, Zhang L, Ren C, et al. Association of CAST gene polymorphisms with carcass and meat quality traits in Yanbian cattle of China. Molecular biology reports. 2013;40(2):1875–1881.

24. Ribeiro VMP, Gouveia GC, de Moraes MM, de Araújo AEM, Raidan FSS, de Souza Fonseca PA, et al. Genes underlying genetic correlation between growth, reproductive and parasite burden traits in beef cattle. Livestock Science. 2021;244:104332.

25. Kern RJ, Lindholm-Perry AK, Freetly HC, Snelling WM, Kern JW, Keele JW, et al. Transcriptome differences in the rumen of beef steers with variation in feed intake and gain. Gene. 2016;586(1):12–26.

26. Keogh K, McKenna C, Porter R, Waters S, Kenny D. Effect of dietary restriction and subsequent realimentation on hepatic oxidative phosphorylation in cattle. Animal. 2021;15(1):100009.

27. Benedeti PDB, Detmann E, Mantovani H, Bonilha S, Serão N, Lopes D, et al. Nellore bulls (Bos taurus indicus) with high residual feed intake have increased the expression of genes involved in oxidative phosphorylation in rumen epithelium. Animal Feed Science and Technology. 2018;235:77–86.

28. Nolte W, Weikard R, Brunner RM, Albrecht E, Hammon HM, Reverter A, et al. Identification and annotation of potential function of regulatory antisense long non-coding RNAs related to feed efficiency in bos taurus bulls. International journal of molecular sciences. 2020;21(9):3292.

29. Hardie L, VandeHaar M, Tempelman R, Weigel K, Armentano L, Wiggans G, et al. The genetic and biological basis of feed efficiency in mid-lactation Holstein dairy cows. Journal of dairy science. 2017;100(11):9061–9075.

30. Lv Y, Cao Y, Gao Y, Yun J, Yu Y, Zhang L, et al. Effect of ACSL3 Expression Levels on Preadipocyte Differentiation in Chinese Red Steppe Cattle. DNA and cell biology. 2019;38(9):945–954.

31. Waters SM, Coyne GS, Kenny DA, Morris DG. Effect of dietary n-3 polyunsaturated fatty acids on transcription factor regulation in the bovine endometrium. Molecular biology reports. 2014;41(5):2745–2755.

32. Li Y, Wang M, Li Q, Gao Y, Li Q, Li J, et al. Transcriptome profiling of longissimus lumborum in Holstein bulls and steers with different beef qualities. PloS one. 2020;15(6):e0235218.

33. Baik M, Vu T, Piao M, Kang H. Association of DNA methylation levels with tissue-specific expression of adipogenic and lipogenic genes in longissimus dorsi muscle of Korean cattle. Asian-Australasian journal of animal sciences. 2014;27(10):1493.

34. Seong J, Yoon H, Kong HS. Identification of microRNA and target gene associated with marbling score in Korean cattle (Hanwoo). Genes & Genomics. 2016;38(6):529–538.

35. Melnik BC, John SM, Schmitz G. Milk consumption during pregnancy increases birth weight, a risk factor for the development of diseases of civilization. Journal of Translational Medicine. 2015;13(1):1–11.

36. Yu SL, Lee SM, Kang MJ, Jeong HJ, Sang BC, Jeon JT, et al. Identification of differentially expressed genes between preadipocytes and adipocytes using affymetrix bovine genome array. Journal of Animal Science and Technology. 2009;51(6):443–452.

37. Engle B, Masters M, Boles JA, Thomson J. Gene Expression and Carcass Traits Are Different between Different Quality Grade Groups in Red-Faced Hereford Steers. Animals. 2021;11(7):1910.

38. Shao T, McCann JC, Shike DW. Effects of Supplements Differing in Fatty Acid Profile to Late Gestational Beef Cows on Steer Progeny Finishing Phase Growth Performance, Carcass Characteristics, and mRNA Expression of Myogenic and Adipogenic Genes. Animals. 2021;11(7):1904.

39. Peletto S, Strillacci M, Capucchio M, Biasibetti E, Modesto P, Acutis P, et al. Genetic basis of Lipomatous Myopathy in Piedmontese beef cattle. Livestock Science. 2017;206:9–16.

40. Martins R, Machado PC, Pinto LFB, Silva MR, Schenkel FS, Brito LF, et al. Genome-wide association study and pathway analysis for fat deposition traits in nellore cattle raised in pasture–based systems. Journal of Animal Breeding and Genetics. 2021;138(3):360–378.

41. de Las Heras-Saldana S, Chung KY, Kim H, Lim D, Gondro C, van der Werf JH. Differential Gene Expression in Longissimus Dorsi Muscle of Hanwoo Steers—New Insight in Genes Involved in Marbling Development at Younger Ages. Genes. 2020;11(11):1381.

42. Zhang F, Wang Y, Mukiibi R, Chen L, Vinsky M, Plastow G, et al. Genetic architecture of quantitative traits in beef cattle revealed by genome wide association studies of imputed whole genome sequence variants: I: Feed efficiency and component traits. BMC genomics. 2020;21(1):1–22.

43. Keogh K, Waters SM, Cormican P, Kelly AK, O’Shea E, Kenny DA. Effect of dietary restriction and subsequent re-alimentation on the transcriptional profile of bovine ruminal epithelium. PloS one. 2017;12(5):e0177852.

44. Srivastava S, Srikanth K, Won S, Son JH, Park JE, Park W, et al. Haplotype-Based Genome-Wide Association Study and Identification of Candidate Genes Associated with Carcass Traits in Hanwoo Cattle. Genes. 2020;11(5):551.

45. Bazile J, Jaffrezic F, Dehais P, Reichstadt M, Klopp C, Laloë D, et al. Molecular signatures of muscle growth and composition deciphered by the meta-analysis of age-related public transcriptomics data. Physiological Genomics. 2020;52(8):322–332.

46. Bernard C, Cassar-Malek I, Le Cunff M, Dubroeucq H, Renand G, Hocquette JF. New indicators of beef sensory quality revealed by expression of specific genes. Journal of Agricultural and Food Chemistry. 2007;55(13):5229–5237.

47. Muniz MMM, Fonseca LFS, dos Santos Silva DB, de Oliveira HR, Baldi F, Chardulo AL, et al. Identification of novel mRNA isoforms associated with meat tenderness using RNA sequencing data in beef cattle. Meat Science. 2020; p. 108378.

48. de Lemos MVA, Peripolli E, Berton MP, Feitosa FLB, Olivieri BF, Stafuzza NB, et al. Association study between copy number variation and beef fatty acid profile of Nellore cattle. Journal of applied genetics. 2018;59(2):203–223.

49. Olivieri BF, Braz CU, Brito Lopes F, Peripolli E, Medeiros de Oliveira Silva R, Ruegger Pereira da Silva Corte R, et al. Differentially expressed genes identified through RNA-seq with extreme values of principal components for beef fatty acid in Nelore cattle. Journal of Animal Breeding and Genetics. 2021;138(1):80–90.

50. Li Y, Wang M, Li Q, Gao Y, Li Q, Li J, et al. Transcriptome profiling of longissimus lumborum in Holstein bulls and steers with different beef qualities. PloS one. 2020;15(6):e0235218.

51. de Almeida Santana MH, Junior GAO, Cesar ASM, Freua MC, da Costa Gomes R, e Silva SdL, et al. Copy number variations and genome-wide associations reveal putative genes and metabolic pathways involved with the feed conversion ratio in beef cattle. Journal of applied genetics. 2016;57(4):495–504.

52. Anton I, Húth B, Füller I, Gábor G, Holló G, Zsolnai A. Effect of single-nucleotide polymorphisms on the breeding value of fertility and breeding value of beef in Hungarian Simmental cattle. Acta Veterinaria Hungarica. 2018;66(2):215–225.

53. Seabury CM, Oldeschulte DL, Saatchi M, Beever JE, Decker JE, Halley YA, et al. Genome-wide association study for feed efficiency and growth traits in US beef cattle. BMC genomics. 2017;18(1):1–25.

54. Manca E, Cesarani A, Gaspa G, Sorbolini S, Macciotta NP, Dimauro C. Use of the Multivariate Discriminant Analysis for Genome-Wide Association Studies in Cattle. Animals. 2020;10(8):1300.

55. Keel BN, Zarek CM, Keele JW, Kuehn LA, Snelling WM, Oliver WT, et al. RNA-Seq meta-analysis identifies genes in skeletal muscle associated with gain and intake across a multi-season study of crossbred beef steers. BMC genomics. 2018;19(1):1–11.

56. Elolimy AA, Moisá SJ, Brennan KM, Smith AC, Graugnard D, Shike DW, et al. Skeletal muscle and liver gene expression profiles in finishing steers supplemented with Amaize. Animal Science Journal. 2018;89(8):1107–1119.

57. Kong RS, Liang G, Chen Y, Stothard P. Transcriptome profiling of the rumen epithelium of beef cattle differing in residual feed intake. BMC genomics. 2016;17(1):1–16.

58. Tizioto P, Coutinho L, Mourão G, Gasparin G, Malagó-Jr W, Bressani F, et al. Variation in myogenic differentiation 1 mRNA abundance is associated with beef tenderness in Nelore cattle. Animal genetics. 2016;47(4):491–494.

59. Leal-Gutiérrez JD, Elzo MA, Johnson DD, Hamblen H, Mateescu RG. Genome wide association and gene enrichment analysis reveal membrane anchoring and structural proteins associated with meat quality in beef. BMC genomics. 2019;20(1):1–18.

60. Ramayo-Caldas Y, Fortes M, Hudson N, Porto-Neto L, Bolormaa S, Barendse W, et al. A marker-derived gene network reveals the regulatory role of PPARGC1A, HNF4G, and FOXP3 in intramuscular fat deposition of beef cattle. Journal of Animal Science. 2014;92(7):2832–2845.

61. Love MI, Huber W, Anders S. Moderated estimation of fold change and dispersion for RNA-seq data with DESeq2. Genome biology. 2014;15(12):1–21.

62. Wheeler T, Shackelford S, Koohmaraie M. Relationship of beef longissimus tenderness classes to tenderness of gluteus medius, semimembranosus, and biceps femoris. Journal of Animal Science. 2000;78(11):2856–2861.

63. Feldsine P, Abeyta C, Andrews WH. AOAC International methods committee guidelines for validation of qualitative and quantitative food microbiological official methods of analysis. Journal of AOAC International. 2002;85(5):1187–1200.

64. Li A, Horvath S. Network neighborhood analysis with the multi-node topological overlap measure. Bioinformatics. 2007;23(2):222–231.

65. Bianchi FM, Grattarola D, Alippi C. Spectral clustering with graph neural networks for graph pooling. In: International Conference on Machine Learning. PMLR; 2020. p. 874–883.

66. Kingma DP, Ba JL. Adam: A method for stochastic gradient descent. In: ICLR: International Conference on Learning Representations; 2015. p. 1–15.

67. Lloyd S. Least squares quantization in PCM. IEEE transactions on information theory. 1982;28(2):129–137.

68. Kaufman L, Rousseeuw PJ. Finding groups in data: an introduction to cluster analysis. vol. 344. John Wiley & Sons; 2009.

69. Yu G, Wang LG, Han Y, He QY. clusterProfiler: an R package for comparing biological themes among gene clusters. Omics: a journal of integrative biology. 2012;16(5):284–287.

